# The Effect of Vaccination on the Evolution of the SARS-CoV-2 B.1.351 Variant

**DOI:** 10.64898/2026.05.06.723356

**Authors:** Ziyi Wang, Madeline Raeihle, Shannon Braun-Gorman, Isaac Leung, Claire Richards, Lea Gabbay, Michel Shamoon-Pour

**Affiliations:** Stony Brook University, Stony Brook, NY 11794; Binghamton University, Vestal, NY 13902; Anthropology Department, Binghamton University, Vestal, NY 13902; First-Year Research Immersion Program, Binghamton University, Vestal, NY 13902

**Keywords:** SARS-CoV-2, Beta Variant, Coronavirus disease, COVID-19, Vaccines, Global Vaccination Rate, Mutation Rate, Nucleotide Diversity

## Abstract

Since the initial distribution of the SARS-CoV-19 vaccine, its widespread use has been hypothesized to act as a selective pressure that drives the COVID-19 virus to mutate. This study aims to investigate the correlation between global vaccination rates and the mutation rate of the SARS-CoV-2 Beta variant (B.1.351). From January to July 2021, nucleotide diversity increased in tandem with vaccination rates, demonstrating that the virus evolved more rapidly in response to selective pressure from mass vaccination. Statistical analysis revealed statistically significant positive correlations between both vaccination rates and vaccine doses administered with nucleotide diversity. Thus, our findings indicate a positive correlation between rising vaccination rates and nucleotide diversity, suggesting that increased vaccination coverage acted as a selective pressure that accelerated viral evolution of SARS CoV2.

## 1. Introduction

### 1.1 Background on SARS-CoV-2

SARS-CoV-2 is a respiratory virus part of the coronaviridae family, and is colloquially referred to as COVID-19 [1]. This virus was initially identified in December 2019 in Wuhan, China, and has subsequently spread worldwide, with 779,207,696 confirmed cases as of March, 2026 [2, 3]. SARS-CoV-2 is classified as the B lineage of *Betacoronavirus*, and it is related to both the SARS and Bat SARS-Like WIV1 coronaviruses [4]. Its genome length is estimated to be 29.8 kB, including six common and six accessory open reading frames (ORFs) [5]. Since its emergence, SARS-CoV-2 has experienced several key mutations within its viral genome, giving rise to multiple genetically distinct variants [6].

The SARS-CoV-2 lineage B.1.351, also known as the Beta variant, was first reported in South Africa in early November 2020 and named as a Variant of Concern (VOC) on December 18, 2020 [6]. The classification of “Variant of Concern” is assigned to variants that may increase in transmissibility and severity, or decrease in neutralization by antibodies, vaccines, or other treatments [7]. The Beta variant was considered a threat to public health due to its unprecedented rate of hospitalization among infected individuals as compared to the previous Alpha variant [8, 9]. In response, multiple vaccines were developed to help induce immunity to the infection and reduce the severity of symptoms.

### 1.2 The Structure of SARS-CoV-2

SARS-CoV-2 is an RNA virus with a positive-sense strand (+ssRNA) genome [10, 11]. The genome is enclosed by the nucleocapsid (N) protein and surrounded by three structural proteins: spike (S), membrane (M), and envelope (E), as well as 16 non-structural proteins [11]. The S protein plays a major role in viral entry into human cells and consists of S1 and S2 subunits. The S1 subunit contains a region called the receptor-binding domain (RBD), the most immunoreactive site on the protein. Due to its role in facilitating infection, as shown in Figure 1, it is possible that structural changes in the S protein lead to variations in SARS-CoV-2 binding affinity to human host cells [12]. The hACE2 receptor, also known as angiotensin-converting enzyme 2, is located on the external surface of many human cells. Unlike other membrane proteins located on the virus, the S protein interacts directly with the hACE2 receptor. The RBD remains hidden until the virus’s spike protein undergoes a conformational change, causing the S protein to take an open position, which allows for fusion with the host cell [11]. Therefore, any changes in the structure of the RBD, particularly in the S protein, may alter its ability to bind to the hACE2 receptor.

**Figure 1.**
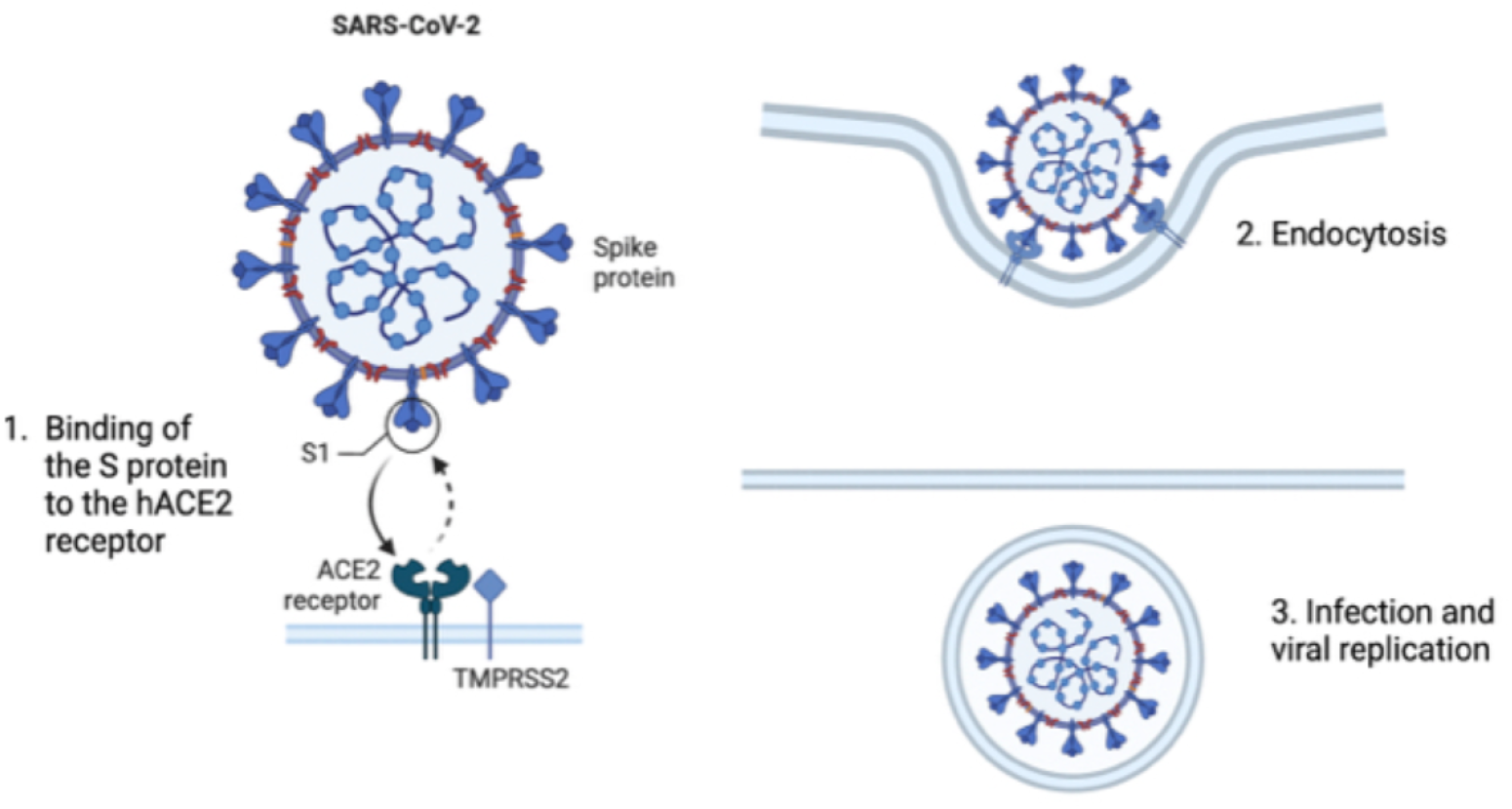
Interaction between SARS-CoV-2 Spike Protein and the hACE2 receptor. 1. The S1 region of the Spike protein attaches to the hACE2 receptor on the cell surface. 2. Endocytosis occurs, and the function of the hACE2 receptor is blocked. 3. Infection occurs, and the viral RNA is replicated within the infected cell.

Certain mutations have been identified as influential in the Beta variant’s increased binding affinity to the hACE2 receptor. Of the 9 mutations on the S protein in the Beta variant, 3 are located on the RBD, and are particularly important to analyze across the sub-lineages of the variant due to the domain’s critical role in the process of infection [8]. However, current data also includes mutations in both the S1 and S2 subunits of the S protein, which are directly involved in antibody evasion [13]. Conserved mutations found in the Beta genome and other variants may be essential to S protein function, as their preservation by natural selection suggests functional importance [14].

### 1.3 The Role of Vaccination

In an attempt to combat the spread of the virus and blunt the severity of its symptoms, multiple vaccinations were developed and distributed. Currently, there are four approved kinds of vaccines: mRNA, inactivated, viral vector, and protein subunit [15]. To evaluate global vaccine rollout, we analyzed vaccination rates in multiple countries (Australia, Brazil, China, Egypt, Greenland, India, Russia, South Africa, Spain, USA) with different authorized vaccines such as Novavax, Moderna, Pfizer, Johnson & Johnson, and Oxford/AstraZeneca. Vaccination rates rose in January 2021 and reached mass coverage (50% population vaccinated) by September 2021 [16]. During this period, the Beta variant was the most widespread VOC, allowing its mutation rate to be easily compared to the percentage of vaccinated individuals.

To investigate external influences on viral evolution, we focused specifically on mutations located on the Beta variant. Mutations on the virus are the result of extrinsic selective pressures, defined as external factors that affect an organism’s evolution [17]. Some proposed examples include mass vaccination and recurrent infections in immunocompromised patients [17]. mRNA vaccines prompt antibody production against the original virus, so widespread vaccination may create selective pressure for SARS-CoV-2 S protein mutations, allowing the virus to evade these antibodies [18]. Additionally, prolonged SARS-CoV-2 infection, especially in hosts who are immunocompromised, may accelerate intrahost viral mutation and support the generation of new variants [19, 20].

### 1.4 Previous studies on vaccination as a selective pressure of SARS-CoV-2

The Red Queen hypothesis suggests that vaccination-driven selection pressures may enhance SARS-CoV-2 transmissibility as an evolutionary arms race arises between SARS-CoV-2 and the human immune system [21]. In this study, a phylogenetic tree was constructed based on the 318 genomes from more than 8500 high-quality SARS-CoV-2 genome sequences in *GISAID* to visualize the major lineages and sub-lineages collected between January 2020 to June 2021 in India. Phylogenetic analysis of the B.1.617 (Delta) variant reveals that the emerging sublineage AY.1 possesses an additional mutation in the RBD, enhancing transmissibility and immune evasion, which likely contributed to its emergence as the dominant strain [21].

### 1.5 Project Objectives and Applications

Because the Beta variant of SARS-CoV-2 remained prevalent globally even amid mass vaccination efforts, its evolution is critical in studying broader mechanisms of viral adaptation as well as the potential effects of mass vaccination. Through analysis of vaccination data from multiple countries and genomic analysis of S protein sequences, we seek to investigate the relationship between mutations in the S protein of the Beta sub-lineages and vaccination rates. Developing an understanding of this relationship may offer valuable insights into future vaccine regulation and development, and assist in informing a more efficient method to control the virus.

## 2. Methodology

### 2.1 Vaccination Data Collection

To assess Beta variant mutation rates in relation to global vaccination coverage, data on monthly vaccination rates from January to December 2021 were extracted from *Our World in Data*, covering the period from the initiation of the vaccine rollout to widespread global uptake. The data were expressed as a percentage of the total population, representing the cumulative worldwide proportion of individuals vaccinated with at least one dose of the SARS-CoV-2 vaccine, as shown in Figure 2.

**Figure 2.**
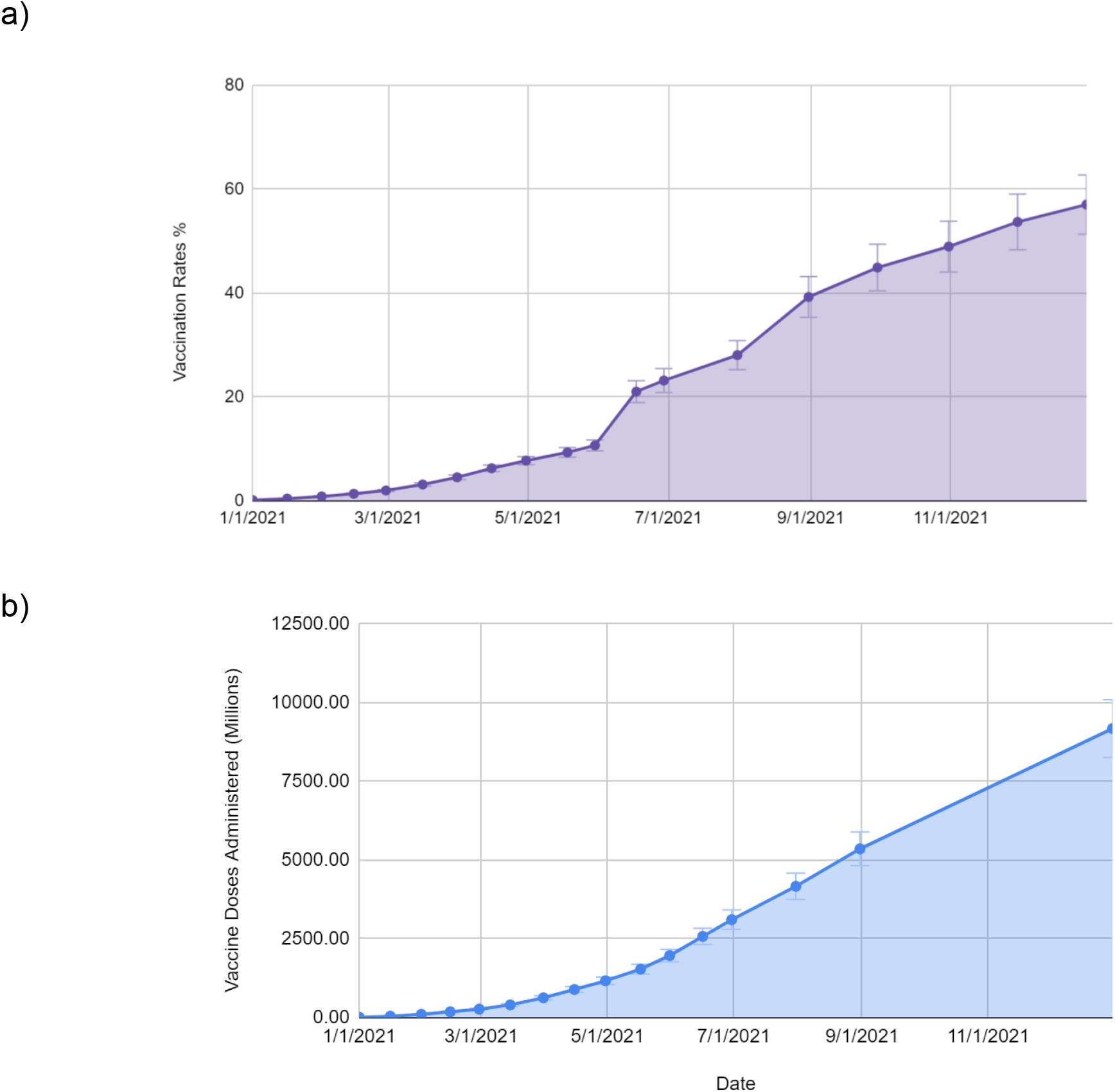
Vaccination data globally through 2021. **a**. Global vaccination trends for SARS-CoV-2 in 2021. The vaccination rate is measured as the percentage of individuals worldwide who have received at least one dose of an SARS-CoV-2 vaccine. **b**. The total amount of SARS-CoV-2 vaccination doses administered throughout 2021 [25].

### 2.2 Sequence Collection and Alignment

To analyze the number of mutations on the SARS-CoV-2 Beta variant, Global 2021 spike glycoprotein sequences were gathered from *GISAID*. The “High-coverage” and “Complete” filters of *GISAID* were used to ensure the validity of the collected sequences. The original Wuhan strain (Wuhan-Hu-1) sequence was used as a reference to compare with the B.1.351 variant, analyzing the evolution and accumulation of mutations over the time period before, during, and after mass vaccination. Through analysis and comparison, we identified the number of mutations in the Spike protein, including insertions, deletions, and substitutions.

*MAFFT* was then used to align the sequences and identify alignment discrepancies [22]. *MEGA 7* was used to manually correct the Spike glycoprotein sequences by removing positions denoted as “-” or “n”, as well as deleting sequences that were missing a large number of positions (∼200) across all sequences [23].

### 2.4 Mutation Analysis

After sequence alignment, the number of mutations in several B.1.351 variant sequences was determined to assess whether they were shared and whether they were affected by SARS-CoV-2 evolution. Tajima’s Neutrality Test in MEGA 11 was used to obtain the nucleotide diversity (π), which measures the average number of nucleotide differences between sequences within each month and reflects genetic variation between strains.

To ensure that nucleotide diversity values were accurate, DnaSP (DNA Sequence Polymorphism) software was used [24]. To read sequence files, ambiguity codes were removed from all the sequences and replaced with “n”. Once the nucleotide diversity values were aligned in MEGA 11 and DnaSP, we can determine whether the π values match across time periods.

The P-value, a statistical measure used to assess whether nucleotide diversity values and vaccination rates are statistically significant or due to chance, was calculated using a one-tailed paired t-test in Google Sheets. The selected ranges represent the nucleotide diversity values and vaccination rates being compared. The tails parameter of the t-test was set to 1, which means a one-tailed distribution and tests for a relationship in one direction, and the type parameter was set to 1 for a paired t-test. We also used the R-test, also known as the correlation test, to assess the strength of the relationship between nucleotide diversity values and vaccination rates. In this analysis, the *R*^2^ value was obtained directly from the graph settings in Google Sheets, and the square-root function was applied to find the Pearson correlation coefficient, r.

## 3. Results and Discussion

### 3.1 Vaccination Rates and Vaccine Doses Administered

Beginning in January 2021, Vaccination rates remained low but started to increase after March 2021. From May 2021 to December 2021, we observed a rapid increase in the global vaccination rate, reaching a threshold of greater than 50%, subsequently attaining mass vaccination status, as shown in Figure 2a. Figure 2b refers to vaccine doses administered between January 2021 and December 2021, and shows a similar pattern in vaccination rate. We observed a sharp rise in the number of individuals with at least one dose of vaccination between June 2021 and July 2021, followed by a significant positive trend from July to December 2021.

### 3.2 Nucleotide Diversity

Figure 3 illustrates nucleotide diversity over time relative to vaccination rates, categorized into three ranges: 0-4.50, 4.60-23.15, and 23.16-67.07. From January 2021 to July 2021, nucleotide diversity increased steadily, reflecting greater genetic variation in Beta sequences over time. However, between August and December, nucleotide diversity declined while vaccination rates continued to rise. This trend may be explained by the decreasing number of recorded Beta variant sequences as the variant declined in August towards the end of the year.

**Figure 3.**
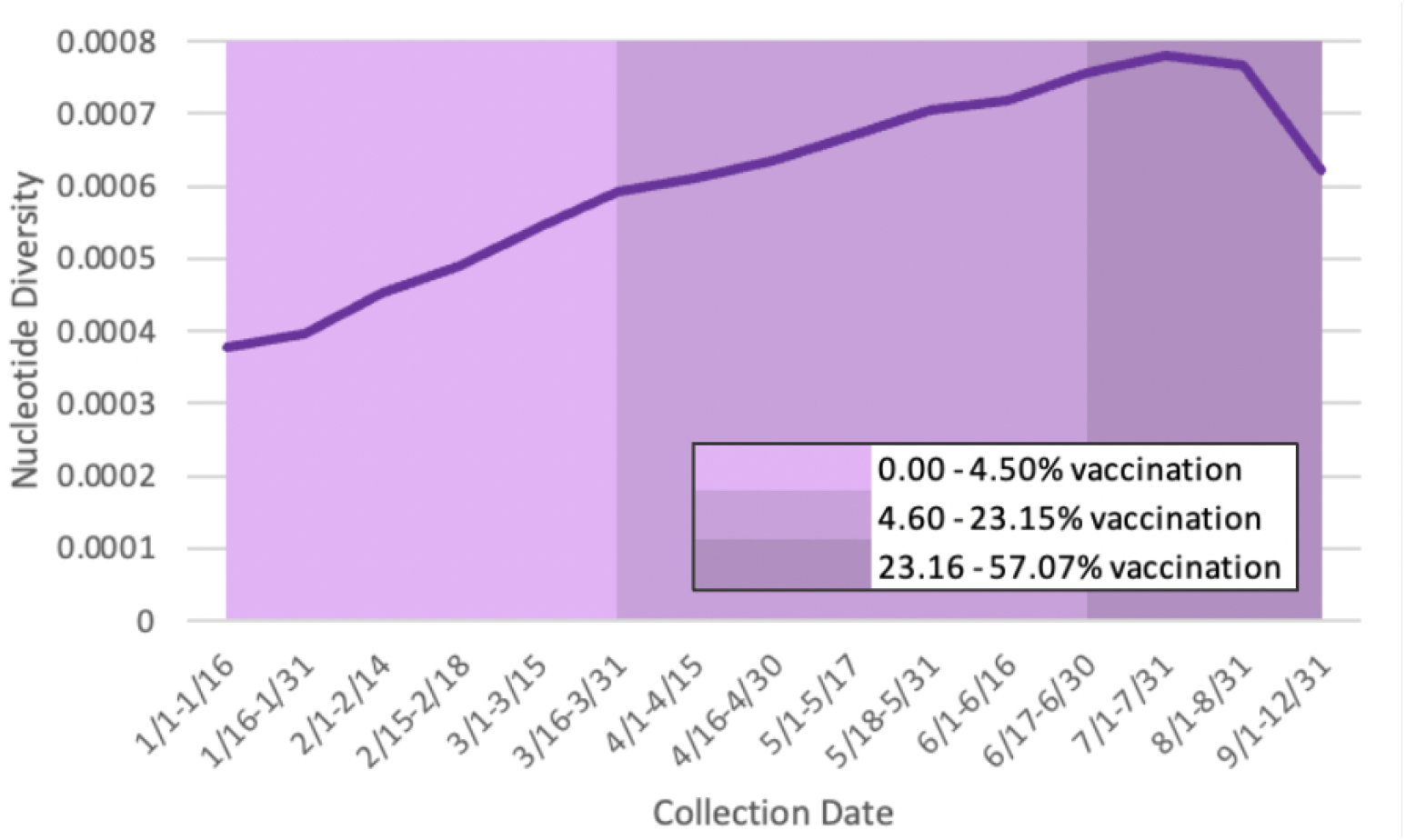
Collection Date vs. Nucleotide Diversity. The nucleotide diversity values for each biweekly group of sequences. This value was calculated by performing Tajima’s Neutrality test in *MEGA11* [26].

### 3.3 Correlation Between Vaccines and Nucleotide Diversity

Data from Figures 2 and 3 were synthesized to create a correlation graph between nucleotide diversity (pi) and vaccination rates (percentage of the world population), as shown in Figure 4a. A line of best fit was included to demonstrate the common trend between nucleotide diversity and vaccination rates. As shown in Figure 4, this indicates a positive correlation, suggesting that as the vaccination rate increased, nucleotide diversity increased as well. The *R*^2^ value of 0.704 in Figure 4a further reinforces the positive association between vaccination rates and nucleotide diversity. Furthermore, the R-value of 0.839 in Figure 4a, which exceeds our significance threshold of 0.5, indicates a strong positive relationship between nucleotide diversity and vaccination rates [27]; the P value of 0.0019 enforces our conclusion of statistical significance.

**Figure 4.**
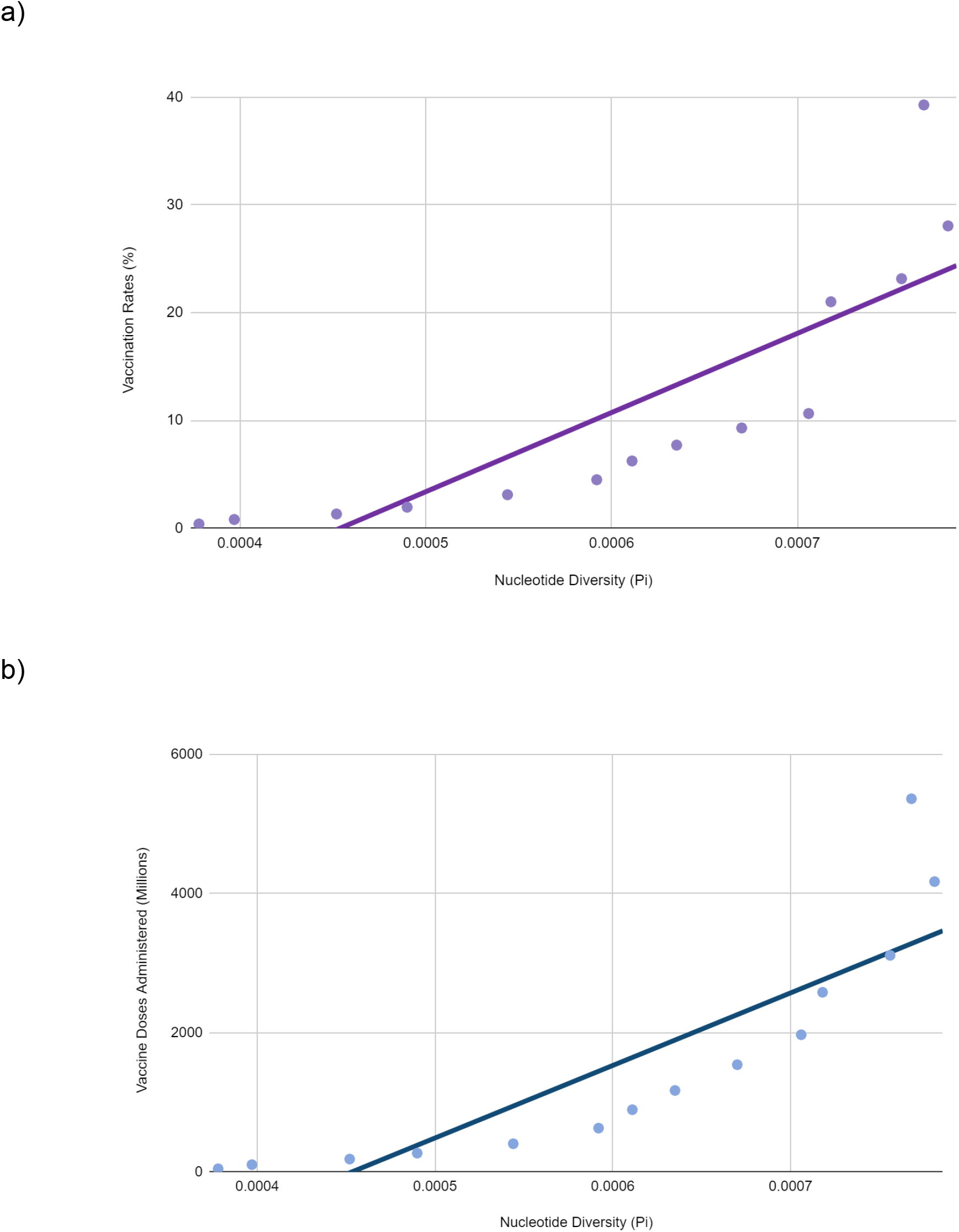
Correlation between Vaccination and Nucleotide Diversity on B.1.351. Figure 4a demonstrates a correlation between vaccination rates, measured as a percentage of the world population, and nucleotide diversity. Figure 4b shows a positive correlation between the number of vaccine doses administered and nucleotide diversity [23, 26, 28, 29].

To investigate the relationship between the number of vaccine doses administered (in millions) and nucleotide diversity (pi), we generated an additional correlation graph. As shown in Figure 4b, the *R*^2^ value of 0.743 indicates a strong alignment of data points to the trendline. 74.3% of the variation in the number of vaccine doses administered can be explained by the Nucleotide Diversity. The R-value of 0.861 further supports a strong positive correlation between nucleotide diversity and vaccine doses administered. The resulting value of 0.0016 from the P-test confirms statistical significance.

Both Figure 4a and 4b demonstrate a strong positive correlation between vaccination rate and nucleotide diversity. In Figure 4a, nucleotide diversity increased alongside rising vaccination rates. Figure 4b illustrates a similar trend between an increase in nucleotide diversity and the number of vaccine doses administered.

Analysis of nucleotide diversity alongside increasing global vaccination rates indicates that vaccination exerted positive selective pressure on the SARS-CoV-2 B.1.351 variant, driving its mutation [30]. While nucleotide diversity declined from August 2021 to December 2021 due to limited sequence data, we observed a significant increase in overall nucleotide diversity as the number of vaccinated individuals increased. These suggest that, within the pairwise sequences collected biweekly, more substitutions and variations were observed as vaccination rates increased throughout 2021. There was also a strong correlation between nucleotide diversity and two measures of global vaccination: the percentage of individuals vaccinated worldwide and the number of vaccine doses administered worldwide. After analyzing both vaccination metrics alongside nucleotide diversity, it is clear that vaccination exerts a positive selective pressure on the virus, favoring the evolution of a resistant strain.

Our conclusion that vaccination is a positive selective pressure on the evolution of SARS-CoV-2 suggests that future studies investigating other epidemiological data may yield more valuable insights. This research also helps policymakers inform decisions on new strategies when facing new SARS-CoV-2 variants. Better safety and hygiene practices are critical to reducing the transmission and contraction of disease. Furthermore, this information may aid in the design and administration of vaccinations to minimize the effect of vaccination as a selective pressure on viral evolution.

### 4. Future Work

While this study demonstrates a correlation between global vaccination rates and the accelerated evolution of the SARS-CoV-2 Beta variant, the findings represent an initial step toward understanding the complex interplay between vaccination and viral adaptation. Due to time constraints and limited data availability, our analysis does not account for the effect of vaccination across countries on the Beta variant’s mutation rate. Additional epidemiological, environmental, and immunological factors may influence viral evolution.

Future research should aim to collect data from diverse global locations to investigate further the environmental factors that may facilitate the evolution of the Beta variant. The high prevalence of the Beta variant within each national population should be considered when selecting specific global locations. Similarities of these locations, such as weather conditions and geographic climate, can be demonstrated. Additionally, comparative data on the effectiveness of different vaccine manufacturers in these high-risk areas could help determine their individual impact on Beta evolution. The integration of additional environmental and immunological studies is necessary to provide further information to identify the new variant and its mutations, and to help local public health authorities respond quickly to new challenges.

By integrating global vaccination data with comparative analyses of vaccine effectiveness, this study helps us better understand what may be driving the mutation and evolution of the SARS-CoV-2 Beta variant. These insights not only clarify the role of vaccination as a potential selective pressure but also highlight the importance of adaptive public health strategies. Ultimately, such findings can optimize future vaccine design and deployment strategies to limit viral evolution and enhance long-term pandemic control more effectively.

## Acknowledgements/Funding

We would like to gratefully thank the First-Year Research Immersion (FRI) program of Binghamton University for supporting this study, as well as Dr. Michel Shamoon-Pour, Vaidik Pandya, and Brad Berrezueta for the mentorship.

## Notes

### Competing Interest Statement

The authors have declared no competing interest.

